# Can we measure engagement from video data based on student movement and sound level?

**DOI:** 10.64898/2026.05.13.723692

**Authors:** Ellen Egeland Flø

**Affiliations:** UiO

**Keywords:** engagement, electrodermal activity, EDA, AI analysis, movement, sound level, quantitative video analysis

## Abstract

Engagement is widely recognised as central to learning and academic achievement. Movement and sound level, combined with an indicator of school relevant content, can be linked to engagement through activating affect, behaviour, and cognition. Electrodermal activity (EDA) is a reliable indicator of movement and sound level, as it measures sympathetic nervous system activation. However, the high cost of wearable EDA sensors has limited its widespread application. This study answers the call for affordable, high-temporal-resolution engagement measures by introducing a video-based quantitative assessment method. 75 minutes of synchronised EDA and video data were collected from 12 upper secondary students (aged 17-18) during regular instruction. Novel software was developed to analyse student movement and sound level for academically relevant content. The OpenPose AI model for pose estimation was also applied. This approach produced six distinct movement variables: two AI-based and four non-AI-based. Six linear models using varying movement variables and sound level were tested to predict tonic EDA levels. All models effectively predicted EDA levels, with non-AI-based movement measures outperforming AI-based alternatives. The four non-AI-based movement models showed similar performance, indicating that compressed versions reduced computational time without sacrificing predictive power. These findings validate a novel, objective method for comparing movement and sound level across learning activities on short timescales, of use for inferring engagement levels. This method is particularly useful for collaborative learning environments and enables controlling for movement and sound in quantitative classroom analyses.

## 1. Introduction

Engagement, a multi-faceted concept encompassing behavioural, affective, and cognitive dimensions (Fredricks et al., 2016; Salas□Pilco et al., 2022), is central for academic performance and learning outcomes (Finn & Zimmer, 2012; Moubayed et al., 2020). However, Macfarlane and Tomlinson (2017) identify issues with less rigorous methods for the measurement of engagement, such as student surveys or teachers’ perceptions of satisfaction. Additionally, using only self-report scales for engagement assessment may introduce bias due to conformity effects (Schnitzler et al., 2021). This bias can be mitigated through real-time measurements, such as video, observations or physiological data (Bae & DeBusk-Lane, 2019; Dubovi, 2022; Li & Lerner, 2013). High-temporal-resolution methods for engagement measurement enable researchers to compare different instructional activities within a single lesson, addressing the call for short-term assessment methods to evaluate interventions aimed at improving student engagement (Bae & DeBusk-Lane, 2019; Li & Lerner, 2013).

These real-time measurements include indicators of physiological arousal, i.e., activation of the sympathetic nervous system (Villanueva et al., 2018). The sympathetic nervous system governs processes that non-consciously regulates the mobilisation of the human body for action (Dawson et al., 2000). Such activation can be measured through several physiological measures e.g., heart rate (variability), respiratory frequency, and electrodermal activity (EDA) (Horvers et al., 2021). Yet the relatively high cost of wearable sensors prevents more widespread use.

However, engagement can be defined on a spectrum from deactivation to activation where high engagement is reflected in activating behaviours, affect, and cognition (Wong & Liem, 2022). Such activation can be observed through increased student movement and sound levels during academically relevant work, where both movement and sensory input increase sympathetic activation (Bari et al., 2023; Christensen & Galbo, 1983; Flø, 2026; Gatti et al., 2018; Lind, 1964; Martin et al., 1974; Weijs et al., 2023). Since EDA is a validated measure of sympathetic activation (Boucsein, 2012), measures of movement and sound level (i.e., sensory input) should predict EDA levels.

Thus, this study aims to validate a novel, objective method for comparing movement and sound level across learning activities on short timescales, of use for inferring engagement levels.

## 2. Engagement

Engagement is a multifaceted term (Groccia, 2018; Martin & Borup, 2022) that refers to which degree a student demonstrates active involvement in both learning and social aspects of the school. The three most common dimensions of engagement are behavioural (e.g., student participation, interaction, collaboration, completion of learning activities), affective (e.g., attitudes and feelings towards teachers, peers, and courses), and cognitive (e.g., motivation and effort to learn) engagement (Fredricks et al., 2016; Salas□Pilco et al., 2022).

A positive relationship exists between student engagement and academic performance (Finn & Zimmer, 2012; Kuhlmann et al., 2024; Moubayed et al., 2020; Zhao et al., 2024), and student engagement is central for learning and decreasing drop-out risk (Archambault et al., 2022; Engels et al., 2021; Hascher & Hagenauer, 2010; Li et al., 2020; Ray et al., 2020; Wang et al., 2017). Moreover, engagement is malleable by structural or large-scale pedagogical development efforts (Finn & Zimmer, 2012). On shorter timescales, engagement can also be affected by teachers’ pedagogical decisions, practice, and the presentation of subject content (Flø & Zambrana, in revision; Finn & Zimmer, 2012).

Engagement needs to be reliably and validly measured to study whether different learning activities improve engagement. Moubayed and colleagues (2020) did this using an unsupervised machine learning model with metrics related to the frequency of student interaction with the course materials and completion of course tasks in a digital environment. Their goal was to model clusters of students based on their engagement levels. Still, no validation of the measure against other measures was sought (i.e., unsupervised machine learning), raising concerns about its validity. On the other hand, D’Mello and colleagues (2017) describe 15 studies that validate their engagement measures. However, these studies’ engagement measurements were confined to digital environments rather than a general classroom setting, which the current study seeks to address.

Another concern regarding measuring engagement is connected to the level at which it is modelled. Differences in investigated student engagement are mainly situated at the student level, i.e., the variance in student engagement is due to differences between students (de Bilde et al., 2013). Consequently, when student engagement is traditionally assessed using self-report measures for models that include individual outcomes, such as academic achievement or school retention, the variance is modelled at the individual level with low temporal resolution (i.e., typically before and after an intervention spanning one or multiple lessons).

Such an approach may pose a challenge for intervention studies aiming to improve student engagement through lesson design interventions, particularly when using several learning activities during lessons. The quantitative video analysis method proposed in the current study will enable this type of analysis, as it measures student engagement across the whole class at high temporal resolution, allowing testing of engagement levels for different learning activities within lessons.

### 2.1 Engagement in the current study

This study adopts a multicomponent perspective on engagement, aligning with the Dual Component Framework of Student Engagement (Wong & Liem, 2022), which synthesises several prior models of engagement into a single framework. Wong & Liem (2022) describe student engagement as comprising learning engagement and school engagement, where learning engagement builds on the common factors of affective, behavioural, and cognitive engagement. The affective factor is described along a spectrum of activation towards deactivation, where engagement is characterised by activating affect (e.g., feeling alert, vigorous, interested), and disengagement by deactivating affect (e.g., feeling bored, tired). The behavioural factor ranges from exertion to withdrawal, and such engagement is marked by intentional exertion of effort (e.g., sustained effort, diligence), and disengagement as intentional withdrawal of effort (e.g., giving up). Lastly, the cognitive factor ranges from absorption to distraction, with engagement reflected in concentration and task-relevant thoughts or processing (e.g., attention), and disengagement reflected in distraction and off-task thoughts (e.g., mind wandering).

The operationalisation in the current study pools the activating affect, intentional exertion of effort, and cognitive absorption. Activating affect is associated with increased movement and sound levels in students compared to deactivating affect, when students feel bored or tired. Behavioural exertion of effort is also associated with increased movement and sound levels in students, compared to behavioural withdrawal, where students give up, as long as the giving up is not associated with extra-curricular engagement. Extra-curricular engagement is here defined as when a student is very enthusiastic, gestures a lot, and talks loudly about something non-academic. To control for such instances and include cognitive engagement, the current study employs a qualitative assessment of academic content based on students’ oral production of academic content.

More specifically, the quantitative video analysis is based on the following definition of learning engagement, which consists of two overlapping factors:

1. Emotional engagement: students’ level of activating affect, e.g., body language, tone and intensity of voice, and the content of their vocal expressions. This is operationalised as higher activating affect, reflected in increased movement and sound levels, when academic content is present.
2. Bodily engagement: students’ level of activity tied to their subject content, e.g., by analysing their amount of movement and their sound level. This is operationalised as increased activity, reflected in higher movement and sound levels, when academic content is present.

These factors draw on the classical behavioural, affective, and cognitive dimensions of engagement (Fredricks et al., 2016; Salas□Pilco et al., 2022) as well as the Dual Component Framework of Student Engagement (Wong & Liem, 2022). Figure 1 shows the study’s definition of engagement. This operationalisation is suitable for assessment on shorter timescales and allows comparison of different learning activities in terms of engagement.

**Figure 1.**
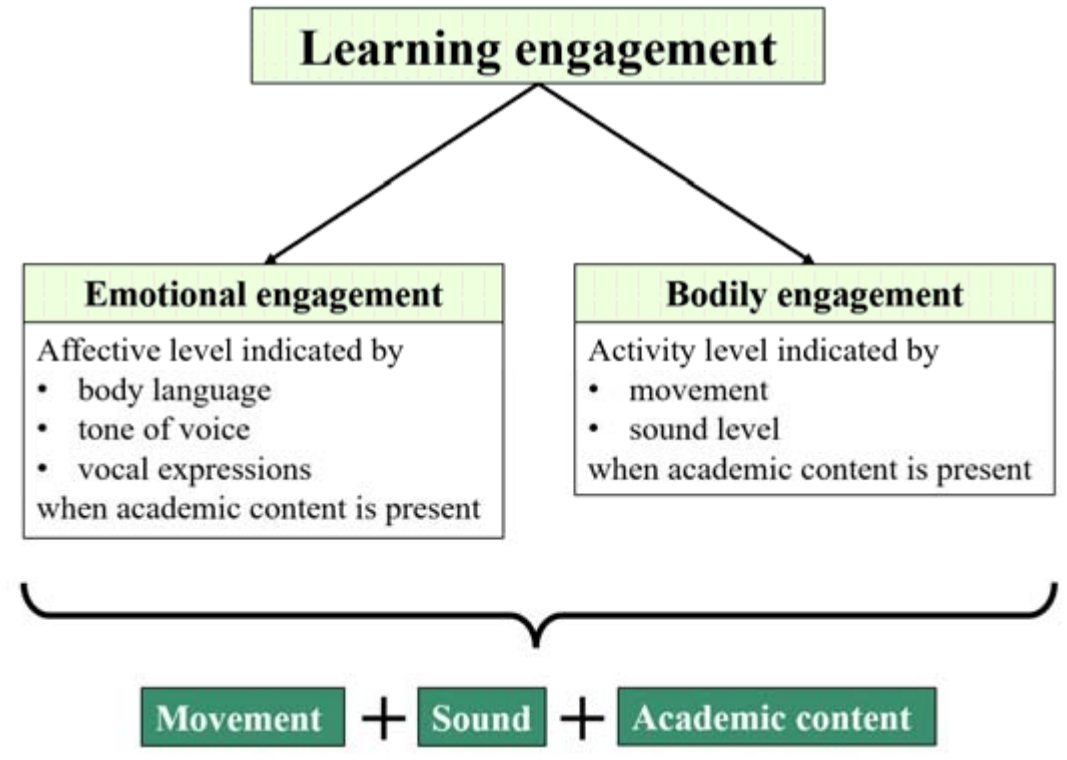
An overview of the definition and operationalisation of learning engagement in the current study.

Given the link between engagement, learning and achievement (Finn & Zimmer, 2012; Moubayed et al., 2020), a gap remains: accessible solutions for engagement measures with high temporal resolution in classroom contexts are lacking (Bae & DeBusk-Lane, 2019; Li & Lerner, 2013; Wong & Liem, 2022). The current study aims to fill this gap in methodological engagement research. Learning engagement can be operationalised as a combination of student movement and sound level when academic content is present. Movement and sound level in segments with academic content are therefore compared to sympathetic activation, which is affected by movement and sensory input (Bari et al., 2023; Christensen & Galbo, 1983; Flø, 2026; Gatti et al., 2018; Lind, 1964; Martin et al., 1974; Weijs et al., 2023). This is done by addressing the following research question:

RQ: Are sound intensity and movement measured from video data related to learning engagement as measured by sympathetic activation?

More specifically, this study examines the predictive power of sound intensity and movement for sympathetic activation (operationalised via EDA) across three video-processing approaches. First, a model using uncompressed video data establishes baseline predictive relationships. Second, three equivalent models with compressed video data evaluate whether compression strategies maintain predictive efficacy while reducing computational demands and quantify the resulting accuracy-efficiency trade-off. Third, two AI-processed video models isolate individual movement while controlling for extraneous object motion. The final model additionally normalises movement by each individual’s estimated area to calculate their weighted contribution to EDA outcomes. This three-tiered approach systematically evaluates computational efficiency, predictive accuracy, and methodological refinement.

## 3. Method

### 3.1 Participants and context

The data were collected in a physics classroom in vocational education, with 12 male students (17-18 years old) and one teacher. EDA was collected for all students, as was the video data. Video data were collected from one camera located at the front of the classroom, facing the students. The data was collected during one 135-minute lesson, excluding two 10-minute breaks. The teacher went through several physics tasks that the students had worked on previously, whilst interacting with them. Afterwards, the students worked further on the same or similar physics tasks, while the teacher helped them. Most of the lesson time, the students were active and academically verbal, interacting with the teacher and each other.

The study was approved by the Norwegian Centre for Research Data (ref. 489494). In accordance with the ethical guidelines of the National Committee for Research Ethics in the Social Sciences and the Humanities (2022), written informed consent from the teacher and students were obtained via a digital portal. If not all students had agreed to the video data collection, the camera would have been angled so that the students who did not agree would be outside the camera’s field of view.

### 3.2 Electrodermal activity data collection and analysis

Electrodermal activity measurements are the most common measure of sympathetic activation in learning contexts due to their relatively low cost and ease of use (Horvers et al., 2021). They measure how well the skin conducts a current when a low voltage is applied (Dawson et al., 2000). This conductivity of the skin is connected to activation of the sympathetic nervous system through the eccrine sweat glands, most abundantly found in the palms and feet, which emit more sweat when aroused (i.e., the sympathetic nervous system is activated). Because sweat consists of water and electrolytes, it acts as an electrical conductor, increasing the skin’s conductivity when sympathetic activation increases.

Due to validity concerns with wearable EDA instruments (Hu et al., 2024; Ronca et al., 2023), the BITalino EDA sensor was employed, which has been validated against laboratory measures (Batista et al., 2019). EDA was recorded at 10 Hz using two solid-gel self-adhesive disposable Ag/AgCl electrodes attached to the second and third proximal phalanges of the non-dominant hand, exceeding the 5 Hz minimum sampling rate required for spontaneous EDA responses (Tronstad et al., 2022). Classroom temperature was maintained at 22°C to eliminate potential thermoregulatory interference with EDA measurement (Boucsein, 2012), consistent with the school’s standard thermostat setting.

The MATLAB based software Ledalab (version 3.4.9; http://www.ledalab.de/) was used for EDA signal processing. The raw signal is the default input to the Ledalab software for decomposition analysis of the EDA signal (Benedek & Kaernbach, 2010), and no preprocessing of the EDA signal was necessary due to excellent data quality, based on visual inspection. Continuous decomposition analysis was employed to decompose the EDA signal into tonic (slow-varying) and phasic (fast-varying) components, as this is recommended for such analyses (Benedek & Kaernbach, 2010). The focus here is on the tonic component of the EDA signal, as the fast-varying phasic component is considered a more immediate and reactive response and has often been linked to cognitive load (Pijeira-Díaz et al., 2018).

Because EDA does not respond instantly but has a delay (latency) of at least one second after stimulus (Sjouwerman & Lonsdorf, 2019; Society for Psychophysiological Research Ad Hoc Committee on Electrodermal Measures, 2012), the analyses used one-minute data segments. Figure 2 provides an example of an EDA signal decomposed into the slow-varying tonic component and the fast-varying phasic component.

**Figure 2.**
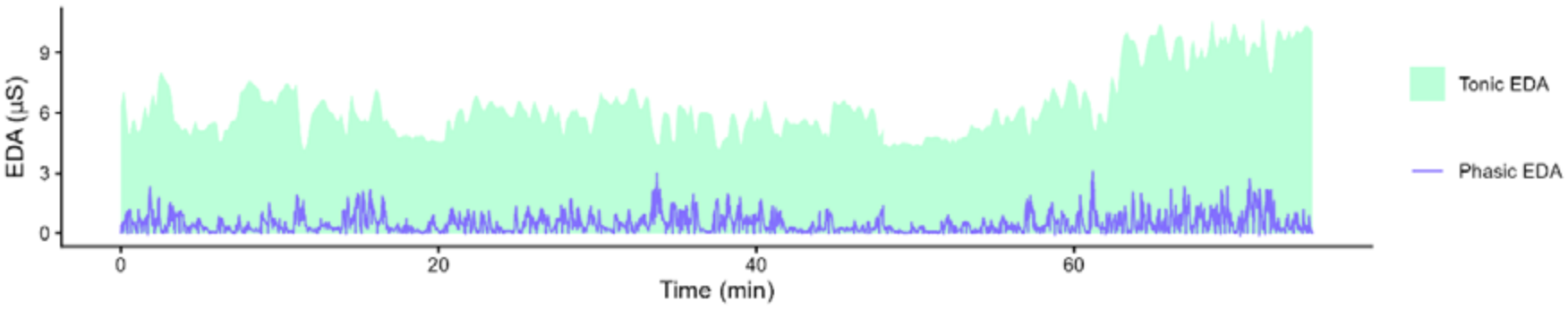
An example of the decomposed EDA signal for one student, where the tonic and phasic components have been separated. The full EDA signal consists of the sum of the tonic and phasic components.

### 3.3 Video data collection and analysis

The quantitative analysis of video data produced mean values per minute for normalised student movement and sound levels. To minimise interference from camera movement and zoom, all video data was collected without zoom from a stationary camera. The camera placement and angle covered almost the entire classroom, excluding only the very front, thereby never leaving any student out of the frame. The practicalities of fastening the EDA sensors, setting up the camera, and disabling EDA collection during breaks yielded 75 minutes of video data with 12 students in an authentic classroom setting.

#### 3.3.1 Qualitative assessment of school content

To ensure that school related engagement is assessed, the video data must be screened for school-related content. In this particular case, the lesson was in physics. Thus, minute segments containing physics content were included in the quantitative analysis, encompassing all 75 available segments. Physics content was assessed independently and to complete agreement by the author and a research assistant, who are both trained in physics at university level, by the presence of spoken physics concepts, units, or related mathematical concepts such as:

specific heat capacity, kilo, degree, formula, heat, mass, kilojoule, strain, elastic, multiply, divide, megapascal, prefix, Young’s modulus, tensile stress, compressive stress, neutron radiation, electromagnetic radiation, light, photocells, photon, wave, particle, diode, light, radio radiation, infrared radiation, ultraviolet radiation, gamma radiation, spectrum, wavelength, electron, proton, sound, vacuum, constructive interference, destructive interference, amplify, frequency, velocity, effect, measure, decibel, sound intensity, Watt, metre, and unit.

This approach was deemed suitable to the data as the class was very vocal and most of the lesson included whole class discussions. Were that not the case, other more non-verbal indicators of academic content should be included.

#### 3.3.2 Uncompressed movement and sound calculations

Custom-developed software (GitHub-link removed for anonymised review) was used to calculate movement. Within each one-minute video segment, the difference between the red, green, and blue (RGB) values of each pixel and the corresponding RGB values in two consecutive frames was calculated. To account for varying lighting conditions, each pixel change value was normalised by dividing it by the corresponding prior pixel value, yielding a measure of change relative to the background. This process was applied to all pixels across all 1499 paired frames in each one-minute video segment (for 25 Hz video), quantifying movement, i.e., changes in student positions. Equation (1) describes how the software computes normalised movement per minute:

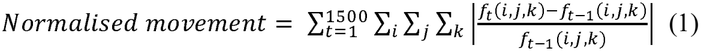

where the function *f* is the individual frame of the video, *i* is the height of the frame, *j* is the width of the frame, *k* is the colour of the pixel, and *t* is the time in 1/25 seconds.

The sound intensity was calculated directly from the video file’s audio (WAV file). The sound intensity was determined by summarising the amplitude of the audio signal for each minute of video. As the sound signal inherently measures intensity relative to a background level of no sound, further normalisation was unnecessary. Equation (2) describes how the software computes sound level per minute:

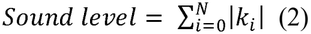

where *k* is the sound level imported from the wav-file and *N* is the number of sound measurements per minute.

#### 3.3.3 Compression in movement calculations

To assess whether it was possible to reliably calculate student movement more efficiently, the video files were down-sampled by frequency, pixel count, or both. As the sound intensity calculation was performed on a much less complex signal, there was no need to down-sample before analysis.

##### Frequency reduction

The video files were down-sampled from 25 frames per second to one frame per second (i.e., 1 Hz). After this compression, normalised movement was calculated as described above.

##### Pixel reduction

Another compression strategy was employed, namely, to reduce the number of pixels included in the calculation of normalised movement. Here, one pixel out of every 20 pixels in both directions was used (instead of all pixels, i.e., ≈ 8.3M pixels), resulting in ≈ 21K pixel values to be summarised. After this compression, normalised movement was calculated as described above.

##### Frequency reduction and pixel reduction

The final and most severe compression was achieved by combining frequency reduction and pixel reduction concurrently.

#### 3.3.4 AI analysis of movement

Student movement was alternatively calculated by using the OpenPose AI model (https://github.com/CMU-Perceptual-Computing-Lab/openpose) for pose estimation. Here, only 1 Hz compressed video was used due to the long calculation time. However, all pixels were retained for each frame, as this was necessary for the OpenPose AI model to reliably identify each participant. Moreover, results were summarised for one-minute segments to be comparable with the other normalised student movement calculations.

The OpenPose AI model generates one JSON file per frame (60 files per minute in this study), each containing coordinates for 25 body parts of identified individuals. To translate these coordinates into movement data, person tracking was implemented across frames using software that matched the closest person in one frame to the subsequent frame (Hur & Bosch, 2022), conducted within one-minute segments. Subsequently, custom-developed software (GitHub-link removed for anonymised review) calculated distances between corresponding body parts across the 60 frames within each minute segment.

Equation (3) describes how the software computes the AI-based movement per minute:

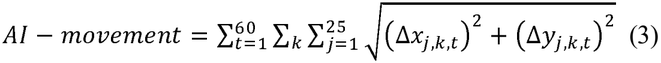

with

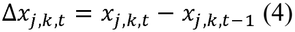

and

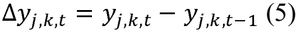

where *j* runs through the 25 key points for the body parts, *k* is each individual person, and *t* is the time in seconds.

Another version of AI-estimated movement, normalised by participant distance from the camera, was calculated. Because the estimated movement will be greater when participants are closer to the camera, as they cover a larger area of the screen, a normalisation that accounted for the area of the identified people was used. This normalisation included dividing each sum of all AI estimated movement for one person per frame by the length between the two shoulder points squared, to account for it being an area effect (i.e., a length squared), and that students were mostly sitting, so their visible area did not consist of their whole bodies. Equation (6) describes how the software computes the normalised AI-movement per minute:

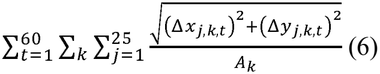

with

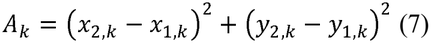

where *j* runs through the 25 key points for the body parts, *k* is each individual person, *t* is the time in seconds, and *A*_*k*_ is the distance between the shoulders for each individual.

Such a normalisation was not possible for the non-AI-based movement calculations as there was no differentiation between individuals.

There was no need to normalise across student number, as all students were present in each frame, which was similarly true for the non-AI-based movement. Thus, the AI-based analysis estimates movement only from people, not from other objects, allowing object movement to be controlled for, unlike the non-AI-based approach. On the other hand, not everyone was consistently identified in each frame for the AI-based movement, particularly when they were sitting at their desks or partly obscured by others. To assess which of these effects impact the tonic EDA level the least, several statistical models were evaluated.

### 3.4 Statistical analysis

R version 4.3.1 (R Core Team, 2023) with R studio version 2023.12.1 was used for the statistical analysis.

First, the pairwise correlations between all variables were calculated using Pearson or Spearman correlation, depending on whether the data were normally distributed, as assessed by the Shapiro-Wilk test. To assess whether sound intensity and movement could predict the tonic component of the EDA signal, and to validate using compressed versions of the movement estimation, four initial multiple regressions were performed with tonic EDA as outcome, with

a. Uncompressed normalised movement and sound intensity
b. Compressed frequency (1 Hz), but with all pixels normalised movement, and sound intensity
c. Non-compressed frequency (25 Hz) with compressed pixels (each 20^th^ pixel in both horizontal and vertical directions) normalised movement, and sound intensity
d. Compressed frequency (1 Hz) with compressed pixels (each 20^th^ pixel in both horizontal and vertical directions) normalised movement, and sound intensity Moreover, two additional multiple regression models were estimated to assess whether AI-based estimates of movement could predict sympathetic activation. Here, tonic EDA was still the outcome, with
e. AI-estimated movement and sound intensity
f. Normalised AI-estimated movement and sound intensity

as regressors. Among these six models (a, b, c, d, e, and f), adjusted R^2^, the Akaike Information Criterion (AIC), the Bayesian Information Criterion (BIC), and Root Mean Square Error (RMSE) were assessed to identify the best fitting model. All regression models were checked for multicollinearity using the variance inflation factor (VIF), linearity using scatterplots and residual plots, homoscedasticity using the non-constant variance test, and normality by visual inspection of the Q-Q plots and the Shapiro-Wilk test.

The resulting best fitting models were compared using a Type III ANCOVA to determine whether they were equivalent by fitting a single combined multiple regression model with one group per model, including interactions between group and sound level, and between group and movement variables.

## 4. Findings

Table 1 presents the pairwise correlations between Tonic EDA, all movement variables, and sound level. All the normalised movements and their corresponding compressed versions demonstrated highly significant correlations, whilst the correlation between the AI-estimated movement variables and the other movement variables were more mixed. Tonic EDA correlated significantly with sound level and one version of normalised movement, although the other normalised movement variables approached significance (p = .087 - .11).

**Table 1.** Pairwise correlation given as Spearman’s rho as the data were not normally distributed, except for AI-based movement and sound level which were both normally distributed, and were thus given as Pearson’s r.

|  | *AI-movement* | *Normalised AI-movement* | *Normalised movement* | *Norm. mov. frequency-compressed* | *Norm. mov. pixel-compressed* | *Norm. mov. full-compressed* | *Sound level* |
|---|---|---|---|---|---|---|---|
| *Tonic EDA* | .08 | -.06 | .20 | .19 | .23* | .20 | .35** |
| *AI-movement* |  | .23* | .40*** | .40*** | .40*** | .40*** | -.15 |
| *Normalised AI-movement* |  |  | -.20 | -.19 | -.24* | -.20 | -.07 |
| *Normalised movement* |  |  |  | .99*** | .98*** | 1*** | -.28* |
| *Norm. mov. frequency-compressed* |  |  |  |  | .98*** | .99*** | -.29* |
| *Norm. mov. pixel-compressed* |  |  |  |  |  | .98*** | -.26* |
| *Norm. mov. full-compressed* |  |  |  |  |  |  | .20 |

\* denotes p-values < .05, \*\* denotes p-values < .01, and \*\*\* denotes p-values < .001

The estimates for the six multiple regression models with tonic EDA as outcome and sound level and one version of movement as independent variables are shown in Table 2. All four normalised movement-based models perform similarly, whilst the AI-based movement estimates show smaller values and lower significance rates.

**Table 2.** Standardised regression coefficients for the multiple regression for tonic EDA data.

| *Model* | *Estimate, β<sub>movement</sub>* | *SE* | *p-value* | *Estimate, β<sub>sound</sub>* | *SE* | *p-value* |
|---|---|---|---|---|---|---|
| *a) Normalised movement* | .34 | .11 | .0021** | .46 | .11 | 4.5e-5*** |
| *b) Norm. mov. frequency-compressed* | .32 | .11 | .0038** | .46 | .11 | 5.4e-5*** |
| *c) Norm. mov. pixel-compressed* | .36 | .10 | .0008*** | .46 | .10 | 3.6e-5*** |
| *d) Norm. mov. full-compressed* | .34 | .11 | .0021** | .46 | .11 | 4.5e-5*** |
| *e) AI-movement* | .16 | .11 | .14 | .40 | .11 | .00044*** |
| *f) Normalised AI-movement* | .084 | .11 | .44 | .38 | .11 | .00083*** |

\* denotes p-values < .05, \*\* denotes p-values < .01, and \*\*\* denotes p-values < .001

Model diagnostics indicated no violations of linearity, homoscedasticity, or normality of residuals. The variance inflation factors were all well below 2.0, suggesting that multicollinearity did not bias the estimates.

Table 3 presents the fit indices for six multiple regression models. Here tonic EDA is the outcome, and sound level and one version of movement are independent variables. All four normalised movement-based models perform similarly well. AI-based movement estimates show poorer fit across all dimensions and the second longest calculation time. Among the four best fitting models, model c) normalised movement pixel-compressed demonstrated the best fit across all dimensions, and a much lower calculation time than uncompressed movement.

**Table 3.** Fit indices for the multiple regression for tonic EDA data.

| *Model* | *R<sup>2</sup><sub>adjusted</sub>* | *AIC* | *BIC* | *RMSE* | *p-value* | *Calculation time/min video* |
|---|---|---|---|---|---|---|
| *a) Normalised movement* | .23 | 198.4 | 207.7 | .861 | 3.5e-5*** | 264 min |
| *b) Norm. mov. frequency-compressed* | .22 | 199.6 | 208.9 | .868 | 6.1e-5*** | 12 min |
| *c) Norm. mov. pixel-compressed* | .25 | 196.6 | 205.9 | .851 | 1.4e-5*** | 15 min |
| *d) Norm. mov. full-compressed* | .23 | 198.4 | 207.7 | .861 | 3.5e-5*** | 1 min |
| *e) AI-movement* | .14 | 206.1 | 215.4 | .907 | .0014** | 60 min |
| *f) Normalised AI-movement* | .13 | 207.8 | 217.1 | .917 | .0031** | 60 min |

\* denotes p-values < .05, \*\* denotes p-values < .01, and \*\*\* denotes p-values < .001

Lastly, a Type III ANCOVA was conducted to test if the four best-fitting models (a, b, c, and d) differed significantly. A common model was estimated with four groups, each based on a different normalised movement variable. The model included possible interactions between these groups and the two main effect variables, namely normalised movement and sound level. The linear model accounted for 22.1% of the total variance (F(11, 288) = 8.72, p = 2.4e-13). There were significant main effects for sound level (F(1, 288) = 18.91, p < .001) and normalised movement (F(1, 288) = 10.25, p = .002). However, the main effect of group (i.e., the different movement variable types) was not significant (F(3, 288) = 0.00, p = 1.00). Additionally, interaction terms involving group were not significant: sound level × group (F(3, 288) = 0.00, p = 1.00) and normalised movement × group (F(3, 288) = 0.03, p = .99). These results suggest that relationships between normalised movement, sound level, and tonic EDA did not differ significantly across the four groups. Thus, the four models using normalised movement or its compressed versions are equivalent.

## 5. Discussion

Here, the research question asking whether sound intensity and movement measured from video data are related to learning engagement, as measured by sympathetic activation, is discussed. More specifically, an initial discussion of study’s findings is followed by a discussion of what it takes to measure *learning engagement*, then the method’s applications in educational research, and, lastly, the limitations and future directions of the research is presented.

### 5.1 Behavioural and physiological measures

Regression analysis confirmed that both engagement indicators (i.e., emotional and bodily engagement), operationalised as sound level and normalised movement when academic content was present, significantly predicted tonic EDA levels. These results aligns with the study’s approach to measuring engagement through observable emotional and bodily dimensions in video data, in line with existing literature linking engagement to sympathetic nervous system activation (Di Lascio et al., 2018; dos Santos Goussain et al., 2023; Kozanitis, 2023; Terriault et al., 2021; Villanueva et al., 2018).

The four non-AI-estimated movement models demonstrated equivalency in performance, eliminating any rationale for selecting alternatives to the fully compressed version, which achieves a substantial reduction in computational time (approximately 260-fold). Conversely, AI-estimated movement models exhibited inferior model fit and required significantly greater processing resources. These limitations may be mitigated in contexts where students are not predominantly seated, as desk-based positioning impairs AI models’ pose estimation capabilities (Vendrow et al., 2023). Future advancements in AI technology for human pose estimation could potentially address these current shortcomings (Vendrow et al., 2023; Yang et al., 2023) and should be an area of future investigation. Consequently, given both superior model performance and computational efficiency, non-AI-estimated movement models represent the preferable approach at present.

### 5.2 Learning engagement

Importantly, this study’s approach incorporates a qualitative assessment of academic content before quantitative analysis, ensuring that the measured engagement is school-relevant. This follows the Dual Component Framework of Student Engagement (Wong & Liem, 2022), which emphasises the need to consider both the behavioural, affective, and cognitive components of engagement in educational contexts. By analysing only segments in which physics concepts were used in a subject-specific context, it was ensured that the measured engagement was academically meaningful rather than merely reflecting general excitement or activity, thereby reflecting cognitive engagement in addition to behavioural and emotional engagement.

This approach addresses a limitation in current engagement research: the ability to distinguish between academically relevant and irrelevant engagement behaviours. Engagement is proposed to be context dependent, being shaped by specific subjects or instructional settings (Appleton et al., 2008; Christenson et al., 2012; Engels et al., 2021; Flunger et al., 2022; Heemskerk & Malmberg, 2020; Kelly & Zhang, 2016; Skinner & Pitzer, 2012). As both time and subject variations in engagement have been identified (Weich et al., 2024), it is likely that academically relevant and irrelevant engagement also differ. The current study addresses this limitation through a mixed-methods approach that incorporates qualitative content analysis prior to quantitative video analysis. This methodology can enable the capture of both movement and vocal expressions while maintaining academic relevance, making it particularly suitable for collaborative learning environments where students freely discuss and interact, rather than in more digital environments (D’Mello et al., 2017). However, its utility may be limited in contexts involving individual silent work or activities where students do not verbalise their thinking processes.

### 5.3 Applications in Educational Research

The quantitative video analysis method may offer some advantages for educational research. By providing a behavioural proxy for engagement that correlates significantly with physiological arousal (specifically, sympathetic activation measured through tonic EDA), this method offers an alternative to self-report engagement measures, which can be subject to response bias and conformity effects (Macfarlane & Tomlinson, 2017; Schnitzler et al., 2021).

A key strength of this method is its high temporal resolution, which allows analysis of movement and sound level on short timescales (e.g., one-minute segments or less) of use for inferring engagement levels. This enables researchers to compare different instructional activities within a single lesson, addressing the call for short-term assessment methods to evaluate interventions aimed at improving student engagement (Bae & DeBusk-Lane, 2019; Li & Lerner, 2013).

This analytical approach offers control for confounding variables in educational research. For instance, in studies employing physiological measures such as electrodermal activity (EDA) or heart rate variability as indicators of sympathetic nervous system activation (Porges, 2025; Boucsein, 2012), controlling for extraneous variables, such as movement and sound levels, is essential. To isolate these factors at the individual level, a video recording system requiring one dedicated camera per participant would be necessary. Given the widespread adoption of personal digital devices among students, such data collection can be implemented using existing hardware, such as student laptops equipped with built-in cameras, thereby reducing logistical barriers while maintaining ecological validity (D’Mello et al., 2017).

A complementary version of this study’s software quantifies environmental sensory stimuli, such as brightness and loudness (Flø & Flø, 2026). These quantified measures can then be correlated with physiological indicators such as EDA, allowing researchers to investigate how environmental factors influence student engagement. This process aligns with established research on sensory processing in educational contexts (e.g., Butera et al., 2020; Ouellet et al., 2021; Samsen-Bronsveld et al., 2022). By systematically measuring both environmental conditions and corresponding physiological responses, researchers can better understand the mechanisms through which sensory environments impact learning engagement.

### 5.4 Limitations and future research

While the findings are promising, several limitations should be acknowledged. The method has only been investigated with 17-18-year-old male students in a physics classroom. Further validation in different age groups, subject areas, and educational contexts is needed. However, to alleviate context-specific concerns, the behavioural measures of movement and sound were correlated with the physiological measure of tonic EDA, providing theoretical grounding that improves generalisability beyond this study’s specific context.

A methodical limitation of the quantitative video analysis is that the method may be influenced by individual differences in typical movement patterns and vocal expressiveness. Research designs should ideally include the same students in all experimental conditions, serving as their own controls. However, randomised group assignment would also reduce the impact of individual differences across all group-difference outcomes.

Another potential limitation of the quantitative video analysis is the qualitative judgement of school vs. non-school engagement, i.e., whether a specific minute-section contains school-relevant content. This judgement can be operationalised in different ways, and in the current study, the most concrete and reproducible operationalisation was to identify whether students used physics concepts.

In observational research investigating naturalistic instruction to identify sources of student engagement, another qualitative coding approach would be beneficial. Rather than using a dichotomous scale, a validated coding manual for video for the specific purpose addressed by the research video observation is recommended, such as the Protocol for Language Arts Teaching Observation (PLATO; Grossman et al, 2013) and the Classroom Assessment Scoring System (CLASS; Pianta et al., 2010).

To validate the video measures as an engagement measure empirically, in addition to the theoretical argument presented in the current study, correlations to other engagement measures should be sought. Future research can further explore how these video-based measures can be integrated with other measures of engagement, such as self-reports, observational coding, or physiological measures beyond EDA. Combining multiple methods could provide a more comprehensive understanding of student engagement across its behavioural, affective, and cognitive dimensions.

## 6. Conclusion

This study demonstrates that quantitative video analysis of movement and sound can serve as a valid proxy for sympathetic activation and, by theoretical extension, student engagement. Coupled with qualitative content analysis to ensure academic relevance, this method enables researchers with a temporally precise tool for investigating the impact of various pedagogical practices and lesson designs on student engagement through movement and sound levels. As such, this study contributes to the educational research toolkit, particularly for intervention studies comparing teaching activities within lessons.

## Data availability

The video dataset generated and analysed during the current study is not publicly available due to ethical restrictions and participant confidentiality agreements, whilst the remaining data is available from the corresponding author upon reasonable request.

## References

Alchieri, L., Alecci, L., Abdalazim, N., & Santini, S. (2023). Recognition of Engagement from Electrodermal Activity Data Across Different Contexts. In Adjunct Proceedings of the 2023 ACM International Joint Conference on Pervasive and Ubiquitous Computing & the 2023 ACM International Symposium on Wearable Computing (pp. 108–112).

Al-Alwani, A. (2016). A combined approach to improve supervised E-learning using multi-sensor student engagement analysis. American Journal of Applied Sciences, 13(12), 1377–1384.

Appleton, J. J., Christenson, S. L., & Furlong, M. J. (2008). Student engagement with school: Critical conceptual and methodological issues of the construct. Psychology in the Schools, 45(5), 369–386.

Archambault, I., Pascal, S., Olivier, E., Dupere, V., Janosz, M., Parent, S., & Pagani, L. S. (2022). Examining the contribution of student anxiety and opposition-defiance to the internal dynamics of affective, Cognitive and Behavioural Engagement in Math. Learning and Instruction, 79, 101593.

Ba, S., & Hu, X. (2023). Measuring emotions in education using wearable devices: A systematic review. Computers & Education, 200, 104797.

Bae, C. L., & DeBusk-Lane, M. (2019). Middle school engagement profiles: Implications for motivation and achievement in science. Learning and Individual Differences, 74, 101753.

Bari, D. S., Rammoo, M. N. S., Aldosky, H. Y., Jaqsi, M. K., & Martinsen, Ø. G. (2023). The five basic human senses evoke electrodermal activity. Sensors, 23(19), 8181.

Batista, D., Plácido da Silva, H., Fred, A., Moreira, C., Reis, M., & Ferreira, H. A. (2019). Benchmarking of the BITalino biomedical toolkit against an established gold standard. Healthcare technology letters, 6(2), 32–36.

Benedek, M., & Kaernbach, C. (2010a). A continuous measure of phasic electrodermal activity. Journal of Neuroscience Methods, 190, 80–91.

Boucsein, W. (2012). Electrodermal activity (2nd ed.). New York, NY:Springer.

Butera, C., Ring, P., Sideris, J., Jayashankar, A., Kilroy, E., Harrison, L., … & Aziz□Zadeh, L. (2020). Impact of sensory processing on school performance outcomes in high functioning individuals with autism spectrum disorder. Mind, Brain, and Education, 14(3), 243–254.

Christensen, N. J., & Galbo, H. (1983). Sympathetic nervous activity during exercise. Annual review of physiology, 45, 139–153.

Cowley, B., Fantato, M., Jennett, C., Ruskov, M., & Ravaja, N. (2014). Learning when serious: Psychophysiological evaluation of a technology-enhanced learning game. Journal of Educational Technology & Society, 17(1), 3–16.

Dawson, M. E., Schell, A. M., & Filion, D. L. (2000). The electrodermal system. In J. T. Cacioppo, L. G. Tassinary, & G. G. Berntson (Eds.), Handbook of Psychophysiology (2nd ed.). Cambridge, UK: Cambridge University Press.

de Bilde, J., Van Damme, J., Lamote, C., & De Fraine, B. (2013). Can alternative education increase children’s early school engagement? A longitudinal study from kindergarten to third grade. School Effectiveness and School Improvement, 24(2), 212–233.

Di Lascio, E., Gashi, S., & Santini, S. (2018). Unobtrusive assessment of students’ emotional engagement during lectures using electrodermal activity sensors. Proceedings of the ACM on Interactive, Mobile, Wearable and Ubiquitous Technologies, 2(3), 1–21.

dos Santos Goussain, B. G. C., de Moura, R. A., Luche, J. R. D., de Souza Andrade, H., Gomes, F. M., & Silva, M. B. (2023). Electrodermal activity as an indicator of student engagement: a comparative study of traditional and active learning environments.

Dubovi, I. (2022). Cognitive and emotional engagement while learning with VR: The perspective of multimodal methodology. Computers & Education, 183, 104495.

D’Mello, S., Dieterle, E., & Duckworth, A. (2017). Advanced, analytic, automated (AAA) measurement of engagement during learning. Educational psychologist, 52(2), 104–123.

Engels, M. C., Spilt, J., Denies, K., & Verschueren, K. (2021). The role of affective teacher-student relationships in adolescents’ school engagement and achievement trajectories. Learning and instruction, 75, 101485.

Finn, J. D., & Zimmer, K. S. (2012). Student engagement: What is it? Why does it matter?. In Handbook of research on student engagement (pp. 97–131). Boston, MA: Springer US.

Flunger, B., Hollmann, L., Hornstra, L., & Murayama, K. (2022). It’s more about a lesson than a domain: Lesson-specific autonomy support, motivation, and engagement in math and a second language. Learning and Instruction, 77, 101500.

Flø. E. E. & Flø, G. M. (2026 preprint). Sympathetic activation of sensory input and learning. bioRxiv 2026.05.01.722216; doi:10.64898/2026.05.01.722216

Flø, E. E. & Zambrana, I. M. (in revision). Identifying design guidelines in STEM making activities based on student engagement, concept use, and opportunities for learning.

Fredricks, J. A., Filsecker, M., & Lawson, M. A. (2016). Student engagement, context, and adjustment: Addressing definitional, measurement, and methodological issues. Learning and Instruction. 43. Special Issue: Student engagement and learning: theoretical and methodological advances)

Gatti, E., Calzolari, E., Maggioni, E., & Obrist, M. (2018). Emotional ratings and skin conductance response to visual, auditory and haptic stimuli. Scientific data, 5(1), 1–12.

Ghiasi, S., Greco, A., Barbieri, R., Scilingo, E. P., & Valenza, G. (2020). Assessing autonomic function from electrodermal activity and heart rate variability during cold-pressor test and emotional challenge. Scientific reports, 10(1), 5406.

Groccia, J. E. (2018). What is student engagement?. New directions for teaching and learning, 2018 (154), 11–20.

Grossman, P., Loeb, S., Cohen, J., & Wyckoff, J. (2013). Measure for measure: The relationship between measures of instructional practice in middle school English language arts and teachers’ value-added scores. American Journal of Education, 119(3), 445–470.

Hascher, T., & Hagenauer, G. (2010). Alienation from school. International journal of educational research, 49(6), 220–232.

Heemskerk, C. H. H. M., & Malmberg, L. E. (2020). Students’ observed engagement in lessons, instructional activities, and learning experiences. Frontline Learning Research, 8(6), 38–58.

Horvers, A., Tombeng, N., Bosse, T., Lazonder, A. W., & Molenaar, I. (2021). Detecting emotions through electrodermal activity in learning contexts: A systematic review. Sensors, 21(23), 7869.

Hu, X., Sgherza, T. R., Nothrup, J. B., Fresco, D. M., Naragon-Gainey, K., & Bylsma, L. M. (2024). From lab to life: Evaluating the reliability and validity of psychophysiological data from wearable devices in laboratory and ambulatory settings. Behavior Research Methods, 1–20.

Hur, P., & Bosch, N. (2022). Tracking individuals in classroom videos via post-processing OpenPose data. In LAK22: 12th international learning analytics and knowledge conference (pp. 465–471).

Kelly, S., & Zhang, Y. (2016). Teacher support and engagement in math and science: Evidence from the high school longitudinal study. The High School Journal, 99(2), 141–165.

Kozanitis, A. (2023). Comparing real time student situational engagement in traditional and active learning classroom using non-invasive electrodermal measurements. In Proceedings IMCIC-International Multi-Conference on Complexity, Informatics and Cybernetics (pp. 133–138).

Kuhlmann, S. L., Plumley, R., Evans, Z., Bernacki, M. L., Greene, J. A., Hogan, K. A., … & Panter, A. (2024). Students’ active cognitive engagement with instructional videos predicts STEM learning. Computers & Education, 216, 105050.

Lee, V. R. (2021). Youth engagement during making: using electrodermal activity data and first-person video to generate evidence-based conjectures. Information and Learning Sciences, 122(3/4), 270–291.

Li, J., Wong, S. C., Yang, X., & Bell, A. (2020). Using feedback to promote student participation in online learning programs: Evidence from a quasi-experimental study. Educational Technology Research and Development, 68, 485–510.

Li, Y., & Lerner, R. M. (2013). Interrelations of behavioral, emotional, and cognitive school engagement in high school students. Journal of youth and adolescence, 42, 20–32.

Lind, A. R. (1964). The circulatory effects of sustained voluntary muscle contraction. Clin. Sci., 24, 229–244.

Macfarlane, B., & Tomlinson, M. (2017). Critiques of student engagement. Higher Education Policy, 30, 5–21.

Martin, C. E., Shaver, J. A., Leon, D. F., Thompson, M. E., Reddy, P. S., & Leonard, J. J. (1974). Autonomic mechanisms in hemodynamic responses to isometric exercise. The Journal of clinical investigation, 54(1), 104–115.

Martin, F., & Borup, J. (2022). Online learner engagement: Conceptual definitions, research themes, and supportive practices. Educational Psychologist, 57(3), 162–177.

Moubayed, A., Injadat, M., Shami, A., & Lutfiyya, H. (2020). Student engagement level in an e-learning environment: Clustering using k-means. American Journal of Distance Education, 34(2), 137–156.

National Committee for Research Ethics in the Social Sciences and Humanities. (2021). Guidelines for research ethics in the social sciences and the humanities. https://www.forskningsetikk.no/en/guidelines/social-sciences-humanities-law-and-theology/guidelines-for-research-ethics-in-the-social-sciences-humanities-law-and-theology/

Ouellet, B., Carreau, E., Dion, V., Rouat, A., Tremblay, E., & Voisin, J. I. (2021). Efficacy of sensory interventions on school participation of children with sensory disorders: A systematic review. American Journal of Lifestyle Medicine, 15(1), 75–83.

Pianta R. C., Hamre B. K., Mintz S. L. (2010). Classroom Assessment Scoring System (CLASS): Upper elementary manual. Charlottesville, VA: Teachstone.

Pijeira-Díaz, H. J., Drachsler, H., Kirschner, P. A., & Järvelä, S. (2018). Profiling sympathetic arousal in a physics course: How active are students?. Journal of Computer Assisted Learning, 34(4), 397–408.

Pijeira-Díaz, H. J., & Channa, F. (2025). Electrodermal Activity in Learning Sciences Research: A Systematic Literature Review. In Proceedings of the 19th International Conference of the Learning Sciences-ICLS 2025, pp. 2871–2873. International Society of the Learning Sciences.

Porges, S. W. (2025). Polyvagal theory: a journey from physiological observation to neural innervation and clinical insight. Frontiers in Behavioral Neuroscience, 19, 1659083.

Ray, A. E., Greene, K., Pristavec, T., Hecht, M. L., Miller-Day, M., & Banerjee, S. C. (2020). Exploring indicators of engagement in online learning as applied to adolescent health prevention: a pilot study of REAL media. Educational Technology Research and Development, 68, 3143–3163.

Reschly, A. L., & Christenson, S. L. (Eds.). (2022). Handbook of research on student engagement (pp. 3–24). New York: Springer.

Ronca, V., Martinez-Levy, A. C., Vozzi, A., Giorgi, A., Aricò, P., Capotorto, R., … & Di Flumeri, G. (2023). Wearable technologies for electrodermal and cardiac activity measurements: a comparison between fitbit sense, empatica E4 and shimmer GSR3+. Sensors, 23(13), 5847.

Salas□Pilco, S. Z., Yang, Y., & Zhang, Z. (2022). Student engagement in online learning in Latin American higher education during the COVID□19 pandemic: A systematic review. British Journal of Educational Technology, 53(3), 593–619.

Samsen-Bronsveld, H. E., van der Ven, S. H., Bogaerts, S., Greven, C. U., & Bakx, A. W. (2022). Sensory processing sensitivity does not moderate the relationship between need satisfaction, motivation and behavioral engagement in primary school students. Personality and Individual Differences, 195, 111678.

Schnitzler, K., Holzberger, D., & Seidel, T. (2021). All better than being disengaged: Student engagement patterns and their relations to academic self-concept and achievement. European Journal of Psychology of Education, 36, 627–652.

Schroeder, N. L., Romine, W. L., & Kemp, S. E. (2023). A scoping review of wrist-worn wearables in education. Computers and Education Open, 5, 100154.

Sjouwerman, R., & Lonsdorf, T. B. (2019). Latency of skin conductance responses across stimulus modalities. Psychophysiology, 56(4), e13307.

Skinner, E. A., & Pitzer, J. R. (2012). Developmental dynamics of student engagement, coping, and everyday resilience. In Handbook of research on student engagement (pp. 21–44). Boston, MA: Springer US.

Society for Psychophysiological Research Ad Hoc Committee on Electrodermal Measures. (2012). Publication recommendations for electrodermal measurements. Psychophysiology, 49(8), 1017–1034.

Sung, G., Bhinder, H., Feng, T., & Schneider, B. (2023). Stressed or engaged? Addressing the mixed significance of physiological activity during constructivist learning. Computers & Education, 199, 104784.

Terriault, P., Kozanitis, A., & Farand, P. (2021). USE OF ELECTRODERMAL WRISTBANDS TO MEASURE STUDENTS’COGNITIVE ENGAGEMENT IN THE CLASSROOM. Proceedings of the Canadian Engineering Education Association (CEEA).

Tronstad, C., Amini, M., Bach, D. R., & Martinsen, Ø. G.(2022). Current trends and opportunities in the methodology of electrodermal activity measurement. Physiological measurement, 43(2), 02TR01.

Vendrow, E., Le, D. T., Cai, J., & Rezatofighi, H. (2023). Jrdb-pose: A large-scale dataset for multi-person pose estimation and tracking. In Proceedings of the IEEE/CVF Conference on Computer Vision and Pattern Recognition (pp. 4811–4820).

Villanueva, I., Campbell, B. D., Raikes, A. C., Jones, S. H., & Putney, L. G. (2018). A multimodal exploration of engineering students’ emotions and electrodermal activity in design activities. Journal of Engineering Education, 107(3), 414–441.

Wang, M. T., Fredricks, J., Ye, F., Hofkens, T., & Linn, J. S. (2017). Conceptualization and assessment of adolescents’ engagement and disengagement in school. European Journal of Psychological Assessment.

Weich, M., Göllner, R., & Stalder, B. E. (2024). Subject and time specificity of students’ cognitive behavioral, and emotional engagement at school. Learning and Individual Differences, 114, 102511.

Weijs, M. L., Jonauskaite, D., Reutimann, R., Mohr, C., & Lenggenhager, B. (2023). Effects of environmental colours in virtual reality: Physiological arousal affected by lightness and hue. Royal Society Open Science, 10(10), 230432.

Wong, Z. Y., & Liem, G. A. D. (2022). Student engagement: Current state of the construct, conceptual refinement, and future research directions. Educational Psychology Review, 34(1), 107–138.

Yang, Z., Zeng, A., Yuan, C., & Li, Y. (2023). Effective whole-body pose estimation with two-stages distillation. In Proceedings of the IEEE/CVF International Conference on Computer Vision (pp. 4210–4220).

Zhao, J. H., Yang, Q. F., Lian, L. W., & Wu, X. Y. (2024). Impact of pre-knowledge and engagement in robot-supported collaborative learning through using the ICAPB model. Computers & Education, 217, 105069.

